# Synthetic Mucus Biomaterials Enable Localized Therapeutic Antibody Delivery in Inflammatory Bowel Disease

**DOI:** 10.1101/2025.08.15.670558

**Authors:** Taj Yeruva, Sydney Yang, Michele Kaluzienski, Rebecca Louisthelmy, Shadin Doski, Katharina Maisel, Gregg A. Duncan

**Author notes:** Correspondence to: Gregg Duncan.

## Abstract

Inflammatory bowel disease (IBD) is a chronic condition characterized by recurrent gastrointestinal inflammation that requires long-term therapeutic intervention. While anti-TNF-α monoclonal antibodies (mAbs) are effective in maintaining remission in IBD, systemic delivery is associated with immunosuppression, poor targeting efficiency, and high cost. To address these limitations, we developed a synthetic mucin-based hydrogel for localized delivery of TNF-α-targeting mAbs. Mucins are heavily glycosylated biopolymers that naturally bind antimicrobial and anti-inflammatory proteins, making them well-suited for local biologic drug delivery at mucosal sites. Synthetic mucin-based hydrogels were formed by crosslinking mucin harvested from porcine small intestine with a 4-arm PEG-thiol and loaded with mAbs to evaluate biocompatibility, antibody release kinetics, and therapeutic efficacy. *In vitro* studies confirmed cytocompatibility of mucin-based hydrogels and demonstrated sustained release of full-length IgG antibodies, with enhanced release under proteolytic conditions simulating the gastrointestinal environment. Moreover, mucin-based hydrogels alone were found to modulate macrophage activation and dampen inflammation in LPS-stimulated macrophages. Treatment of LPS-stimulated macrophages with mAb-loaded hydrogels reduced pro-inflammatory cytokine production and macrophage activation, confirming retention of mAb bioactivity. Compared to antibodies administered in solution, *in vivo* biodistribution studies revealed greater absorption of antibodies when loaded in mucin-based hydrogels and administered via enema in TNBS-induced colitis mice likely due to enhanced adhesion to mucosal epithelium and slowed intestinal clearance. This study demonstrates the potential of mucin-based hydrogels as a platform for local mAb delivery in IBD, enabling targeted immunosuppression while minimizing systemic exposure.

## Introduction

Inflammatory bowel disease (IBD), such as Crohn’s disease and ulcerative colitis, involves chronic immune dysregulation in the gastrointestinal (GI) tract, marked by unpredictable flares that often necessitate ongoing or repeated courses of anti-inflammatory therapy over a patient’s lifetime. ^1,2^ With many IBD patients eventually becoming unresponsive to corticosteroids, monoclonal antibodies (mAbs) have emerged as highly effective biologic therapies for IBD. Specifically, TNF-α neutralizing mAbs have proven successful in maintaining IBD remission.^3,4^ However, high doses of anti-TNF-α mAbs (∼5 mg/kg intravenous route or ∼100 mg subcutaneous route) are typically given on a routine basis (e.g., every 2 or 6 weeks) until patients achieve remission.^5^ As a result of the systemic delivery route and repeated dosing requirements, mAbs used to dampen inflammation also pose a potential risk of immunosuppression which render the patient susceptible to infections.^6,7^ For example, anti-TNF-α therapy doubles the risk of opportunistic infections such as tuberculosis, herpes simplex infection and candidiasis in IBD patients.^8^ Further, systemic administration of mAbs only allows a fraction of the therapeutic payload to reach their intended tissue making these therapies very costly. Therefore, there is a strong rationale for developing platforms that enable localized delivery of biologics in IBD, allowing therapeutic levels to be achieved with lower overall doses.

Several biomaterial systems have been engineered in prior work for therapeutic delivery through both the intrarectal and oral administration routes to address the shortcomings of current IBD treatments.^9^ For example, previous studies from the Karp, Langer, and Traverso labs developed an inflammation-targeted hydrogel capable of adhering to lesions in the colon enabling sustained local corticosteroid (dexamethasone) delivery resulting in effective therapeutic outcomes in mouse models of IBD.^10^ Importantly, these studies also demonstrated an overall reduction in systemic exposure to dexamethasone when locally delivered via enema in a hydrogel format. To address excessive bleeding in the GI tract that exacerbates disease in IBD, a protein-based hydrogel system was developed for local delivery of heparin which both reduced micro thrombosis incidence and inflammation leading to intestinal tissue repair in a murine model of colitis.^11^ Secreted mucins have also been implicated in disease progression in IBD as the lack of mucus production in the gastrointestinal tract in IBD weakens the body’s defenses against opportunistic pathogens.^12–14^ As it stands, current treatment regimens do not directly address mucosal barrier dysfunction in IBD. Motivated by this, prior work has also developed orally administered, drug-free, *in situ* forming “mucus-like” hydrogels to locally coat inflamed regions of the colon which lead to improvements in disease symptoms in a mouse model of IBD.^15^ Thus, these past studies collectively demonstrate biomaterials may be used to locally deliver drugs and may help protect the colon from further damage to promote resolution of IBD.

Building further on these past studies, we developed mucin-based biomaterials for local delivery of mAbs in IBD to limit systemic exposure and reduce dose requirements for improved therapeutic outcomes. In the gut and other mucosal tissues, mucus is naturally enriched with peptides and antibodies to help control pathogen invasion and to prime the immune system.^16^ These natural binding pockets on mucins make it the ideal biomaterials for encapsulation and controlled release of mAbs. Mucin glycoproteins are also immunologically active and their interactions with immune cells such as macrophages, a central player in IBD, may further augment excessive inflammatory responses.^17,18^ In prior work, we have used these mucin-based biomaterials for delivery of antimicrobial peptides and antibiofilm agents to treat *Pseudomonas aeruginosa* infections.^19,20^ To evaluate the potential use of mAb-loaded synthetic mucus gels to treat IBD, we evaluated the anti-inflammatory effect of this formulation against macrophage activation *in vitro* and performed *in vivo* studies to establish feasibility of local mAb delivery using synthetic mucus biomaterials for IBD treatment.

## Materials and Methods

### Preparation of IgG loaded synthetic mucus (SM) gel

To prepare synthetic mucus gels, porcine small intestinal mucin (PSIM) was extracted and partially purified using a previously reported method.^19^ PSIM solution was prepared by dissolving in phosphate buffered saline (PBS) (≤4% w/v) and stirred for 2 hours at room temperature. The PSIM solution was mixed with a solution containing a crosslinking reagent 4-arm thiol terminated PEG (PEG-4SH; 10kDa; Laysan Bio) dissolved in PBS (≤4% w/v) in equal volumes and left for 24 hours to gel.^21^ To prepare Immunoglobulin G (IgG) loaded SM hydrogel (IgG-SM), 0.1% w/v IgG is added to the PSIM solution before stirring. To achieve IgG loading concentrations at or exceeding half the stock IgG concentration, we instead dissolved the PSIM and PEG-4SH in IgG solution directly before mixing.

### Bulk rheology

To confirm successful formation of synthetic mucus gels and to understand how polymer weight percentages influence its mechanical properties, bulk rheological measurements were performed using an ARES G2 rheometer (TA instruments). One mL of preformed SM gels in a flat cylindrical shape prepared as described previously were loaded onto a 25 mm diameter parallel plate at a gap of 1000 μm at 37°C. A humidity chamber was used to prevent solvent evaporation and the hydrogel drying. To determine the linear viscoelastic region of the fully formed gel, a strain sweep measurement was performed at 0.1–10% strain at a frequency of 1 rad/s. To determine the elastic modulus, G’(ω), and viscous modulus, G” (ω), a frequency sweep measurement is conducted within the linear viscoelastic region of the gel, at 10% strain amplitude and angular frequencies from 0.1 to 100 radians/s. To elucidate shear thinning nature of SM hydrogel, flow sweep was performed from a shear rate of 0.01 to 100 s^−1^.

### Fluorescence recovery after photobleaching

A total of 25□μL of 0.1% w/v FITC-labeled IgG (FITC-IgG; Sigma Aldrich) encapsulated in SM hydrogel precursor solution was loaded into custom-made microscopy chambers and allowed to gel overnight at room temperature. Samples were protected from light until imaging. Fluorescence recovery after photobleaching (FRAP) experiments were performed using a Zeiss LSM 800 confocal fluorescence microscope equipped with a 63× water-immersion objective. Prior to photobleaching, 10 baseline images were acquired. A circular region of interest (ROI1) with a diameter of 25□μm was photobleached using a 488 nm laser at 100% intensity. A second circular region (ROI2) of the same diameter was defined as a reference to normalize fluorescence intensity data. Following photobleaching, fluorescence recovery within the ROIs was monitored for 3 minutes using 2% laser intensity, with images captured every 0.828 seconds. Data is collected from three independent hydrogel samples, with at least two FRAP measurements performed per sample and analyzed according to a previously published protocol.^22^ Briefly, fluorescence intensities are normalized using Equation 1. The normalized recovery curves are used to calculate the mobile fraction (defined as the plateau value) and the immobile fraction (1 − mobile fraction). Recovery kinetics are fitted to Equation 2 to determine the half-time of recovery (t□/□), which was then used to calculate the diffusivity of IgG within the SM hydrogel using Equation 3.

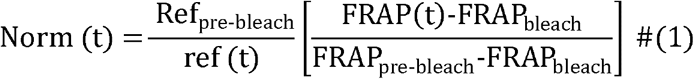

where Ref_pre□ □ □bleach_ is the mean intensity of reference region pre-bleach, FRAP_pre□ □ □bleach_ is the mean intensity of the FRAP region pre-bleach, FRAP_bleach_ is the intensity of the FRAP region at the time of bleaching, ref(*t*) is the intensity of the reference region at time point *t*, and FRAP(*t*) is the intensity of the FRAP region at time point *t*.

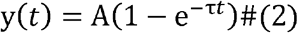

where A equals plateau intensity, _τ_□=□fitted parameter, and *t*□=□time after bleach.

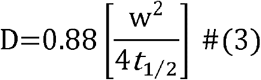

where D is the estimated diffusion coefficient, “*w*” is the bleach area radius, and *t* _1/2_ is the time to reach half the intensity of full recovery.

### Antibody release from SM gels

FITC-IgG was added to the PSIM solution, as described previously, and mixed with equal volumes of PEG-4SH. These solutions were added to tip cut-off syringes, sealed with parafilm, and left undisturbed for 24 h to make 100 μL cylindrical hydrogels. The resulting gels were immersed in 1 mL PBS or PBS with trypsin (143 μg/mL; Worthington Biochemical, LS003740) and pepsin (24.2 μg/mL; Sigma-Aldrich, P700) to simulate the intestinal environment. Samples were incubated at 37 °C, and supernatants were collected at designated time points and replaced with fresh buffer. To examine FITC-IgG release, the fluorescence of the collected supernatants was measured at Ex/Em wavelengths of 490/525 by UV/Vis spectroscopy using a Tecan Spark multimode microplate reader. To determine the mechanism of drug release for cylindrical shaped matrices, first 60% drug release data were fitted in Korsmeyer Peppas model (equation 4).^23^

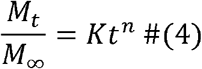

where Mt / M∞ is a fraction of drug released at time t, K is the release rate constant, and n is the release exponent.

### *In vitro* biocompatibility of SM gels

To evaluate the biocompatibility of SM gels *in vitro*, HEK293T cells were treated with SM gels, and cell viability was assessed using a resazurin assay. Briefly, cells were seeded in a 96-well plate at a density of 2□×□10□ cells/well and incubated for approximately 24 hours to reach ∼70% confluence. Cells were then directly treated with 20 μL/well of either unloaded SM gels or SM gels loaded with 0.1% w/v anti-TNF-α monoclonal antibody. Dulbecco’s Modified Eagle Medium (DMEM) served as the negative control, while 1% Triton X-100 was used as the positive control. After 24 hours of incubation, treatment solutions were removed, and cells were gently washed with PBS. Resazurin assay reagent (Biotium) was added to each well following the manufacturer’s protocol and incubated for 3 hours at 37□°C in 5% CO□. Subsequently, 100 μL of the supernatant containing resazurin was transferred to a black 96-well plate, and fluorescence was measured at excitation/emission wavelengths of 570/585 nm using a Tecan Spark multimode microplate reader (5 nm bandwidth for both filters). Cell viability was calculated relative to untreated cells grown in DMEM.

### Immune modulation by mucins and SM gels

Murine RAW 264.7 macrophages (ATCC TIB-71) were cultured in DMEM media with 10% fetal bovine serum and 1% Penicillin-Streptomycin solution at 37°C and 5% CO_2_. Macrophages were seeded at a density of 5×10^5^ per well in tissue culture-treated 24-well plates and incubated overnight at 37°C and 5% CO_2_ to allow the macrophages to attach. Fresh media was added and free PSIM or SM gels were added into the wells and incubated at 37°C and 5% CO_2_ for 24 hours. Untreated macrophages were used as the control. After incubation, the media from the wells was collected and Enzyme-linked immunosorbent assays (ELISAs) were used to measure the concentration of IL-10, IL-6, TNF-α, and TGF-β cytokines produced by RAW 264.7 macrophages. To assess effect of SM hydrogel treatment on pro-inflammatory polarized macrophages, seeded macrophages were stimulated with 10 ng/mL lipopolysaccharide (LPS from *Escherichia coli* O111:B4; Sigma Aldrich, L2630) for 24 hours at 37°C and 5% CO_2_. As described prior, macrophage treatment with PSIM and SM gels was conducted on LPS stimulated macrophages. Media collection was repeated and the concentration of cytokines produced by pro-inflammatory macrophages was measured by ELISAs. Further, expression of anti-inflammatory macrophage cell surface marker (CD 80), pro-inflammatory macrophage cell surface marker (CD 206), and general macrophage marker (F4/80) were measured to assess effect of treatment on macrophage polarization. Phycoerythrin (PE)-anti-mouse CD 80 antibody, allophycocyanin (APC)-anti-mouse CD 206 antibody, Pacific Blue-anti-mouse F4/80, Zombie NIR, and Trustain FcX PLUS antibody were used for staining and all purchased from Biolegend. Staining was performed for both unstimulated and LPS stimulated macrophages treated with PSIM and SM gels. After treatment in 24-well plates, the macrophages were scraped with mini cell scrapers and suspended in PBS. Suspended cells were stained with Zombie NIR in a 500 μL volume, blocked with Trustain FcX PLUS, and incubated at 4°C for 30 minutes. The cell suspension was then stained with the chosen antibodies in cell staining buffer and incubated at 4°C for 60 minutes. The cells were fixed in 4% paraformaldehyde, washed with cell staining buffer, and resuspended in 300 μL of fresh cell staining buffer for analysis using flow cytometry.

### Phagocytosis of pHrodo bioparticles by macrophages

SM hydrogels were added into the 24 well plates seeded with macrophages as described earlier. Untreated macrophages were used as the control. Phagocytosis of pHrodo Red *Escherichia coli* Bioparticles (Thermofisher Scientific) was assessed for macrophages that were unstimulated or stimulated with LPS for 24 hours prior to SM hydrogel treatment. To evaluate the phagocytic activity of macrophages, bioparticles were added at a final concentration of 100 μg/mL per well and were incubated at 37°C and 5% CO_2_ for 2 hours. Wells were then washed with PBS to remove bioparticles not uptaken. To prepare for flow cytometry, macrophages were scraped from the wells, stained with Zombie NIR, and placed on ice until flow cytometry was run. The 488 nm and 640 nm lasers with the PE and APC-Cy7 bandpass filters were used to detect pHrodo red and Zombie NIR, respectively. To image uptake, sterile 14 mm circular glass cover slides were placed into each well of a 24-well plate. Macrophages were seeded, incubated, and treated with SM hydrogels and bioparticles as described prior. After treatment, macrophages were fixed with 1% paraformaldehyde and stained with Alexa Fluor 647 phalloidin (Abcam) and DAPI (Thermofisher Scientific) to visualize the actin filaments and nuclei, respectively. Using sterile tweezers, the circular glass cover slides were removed from the wells and mounted onto rectangular glass slides using Vectashield Antifade Mounting Medium (Vector Laboratories). Images were taken using a confocal fluorescence microscope (Zeiss Confocal LSM 800) fitted with a 25x water-immersion objective. Phalloidin, DAPI, and pHrodo red were detected using 555 nm, 385 nm, and 475 nm lasers, respectively. Four images were taken per well.

### *In vitro* efficacy of anti TNF-α mAb-loaded SM gels

The anti-inflammatory effect of TNF-α targeting mAb-loaded hydrogels was evaluated in RAW264.7 mouse macrophages stimulated with LPS. Briefly, cells were seeded in a 24-well plate at a density of 2□×□10^5^ cells/well and incubated for approximately 24 hours to reach ∼70% confluence. Cells were then stimulated with 100 ng/mL LPS for 24 hours. LPS stimulated macrophages were then treated with 50 μL/well of either unloaded SM gels, SM gels loaded with 0.1% w/v anti-TNF-α monoclonal antibody, or a solution of 0.1% w/v anti-TNF-α monoclonal antibody. Dulbecco’s Modified Eagle Medium (DMEM) served as the negative control. After 24 hours of incubation, treatment solutions were stored for cytokine analysis, and cells were gently washed with PBS. Following washing, macrophage polarization and cytokine production (e.g., TNF-α, IL-6, IL-10) was assessed via flow cytometry and ELISA, as described previously.

### Retention of SM gels following intrarectal delivery

All animal experiments were conducted in accordance with protocols approved by the Institutional Animal Care and Use Committee (IACUC protocol # R-NOV-22-50) at the University of Maryland. Female C57BL/6 mice (6-8 weeks old, The Jackson Laboratory) were used for the study. Mice were fully anesthetized using isoflurane and held vertically by their tails during all administrations. To reduce the passage of pellets prior to trinitrobenzene sulfonic acid (TNBS)-mediated induction of colitis, mice were given a 200□μL saline enema twice using a wiretrol to clear the distal 3–4□cm of the colorectum with a 10 min wait time in between. 10□min post second enema, 65□μL of TNBS (2.5% TNBS in 50% ethanol) was administered intrarectally using a wiretrol (Day 0).^24^ Mice were monitored daily for weight loss until Day 3 to confirm onset of disease. 0.1% w/v Cy5 labeled IgG (Cy5–IgG; Protein Mods) loaded SM gel (excitation: 650 nm, emission: 670 nm) containing 3% w/v PSIM and 2% w/v PEG-4SH was prepared as described previously. On Day 3, mice were given SM gel control (n=2), Cy5-IgG solution (n=3), and Cy5-IgG loaded SM gel (n=5) intrarectally to evaluate retention following saline enema. At 15 minutes and 2 hours post administration, mice were euthanized, and colons were harvested. Fluorescence signals from the Cy5–IgG in the isolated colons were measured using IVIS Spectrum Imaging System. Background signal was recorded from SM gel control mice.

## Results and Discussion

### Formulation and characterization of SM gels for mAb delivery

As shown in **Figure 1a**, we successfully formulated synthetic mucus (SM) gels through the crosslinking of porcine small intestinal mucin (PSIM) with thiolated 4-arm polyethylene glycol (PEG-4SH). We hypothesized substantial loading and extended release of full-length IgG antibodies could be achieved via Fc-mediated interactions with mucin biopolymers in these gels. A range of SM formulations were tested by varying the concentrations of PSIM and PEG-4SH. Some tested formulations either degraded within 8 hours immersed in PBS or were too mechanically stiff for administration via injection (Table S1). Among the tested formulations, the 3% PSIM + 2% PEG formulation was selected as the lead candidate. This formulation could be administered through an 18G needle while maintaining structural stability in PBS for at least 24 hours, making it suitable for controlled drug delivery. The 3% PSIM + 2% PEG SM gels exhibited viscoelastic behavior, with a storage modulus of approximately 45□Pa (**Fig. 1b**). Continuous flow rheology measurements revealed that the viscosity of the SM gels decreased with increasing shear rate, demonstrating shear-thinning behavior (**Fig. 1c**). This property is critical for both injectability and effective spreading at the application site. To evaluate biocompatibility, HEK293T cells were treated with either unloaded SM gels or anti-TNF-α-loaded SM gels. Both treatments maintained over 80% cell viability, in contrast to the positive control (1% Triton X-100), which reduced viability to below 2%. No statistical significance was observed in cell viability between the untreated control group and the SM-treated groups, confirming the biocompatibility of the SM gels (**Fig. 1d**).

**Figure 1.**
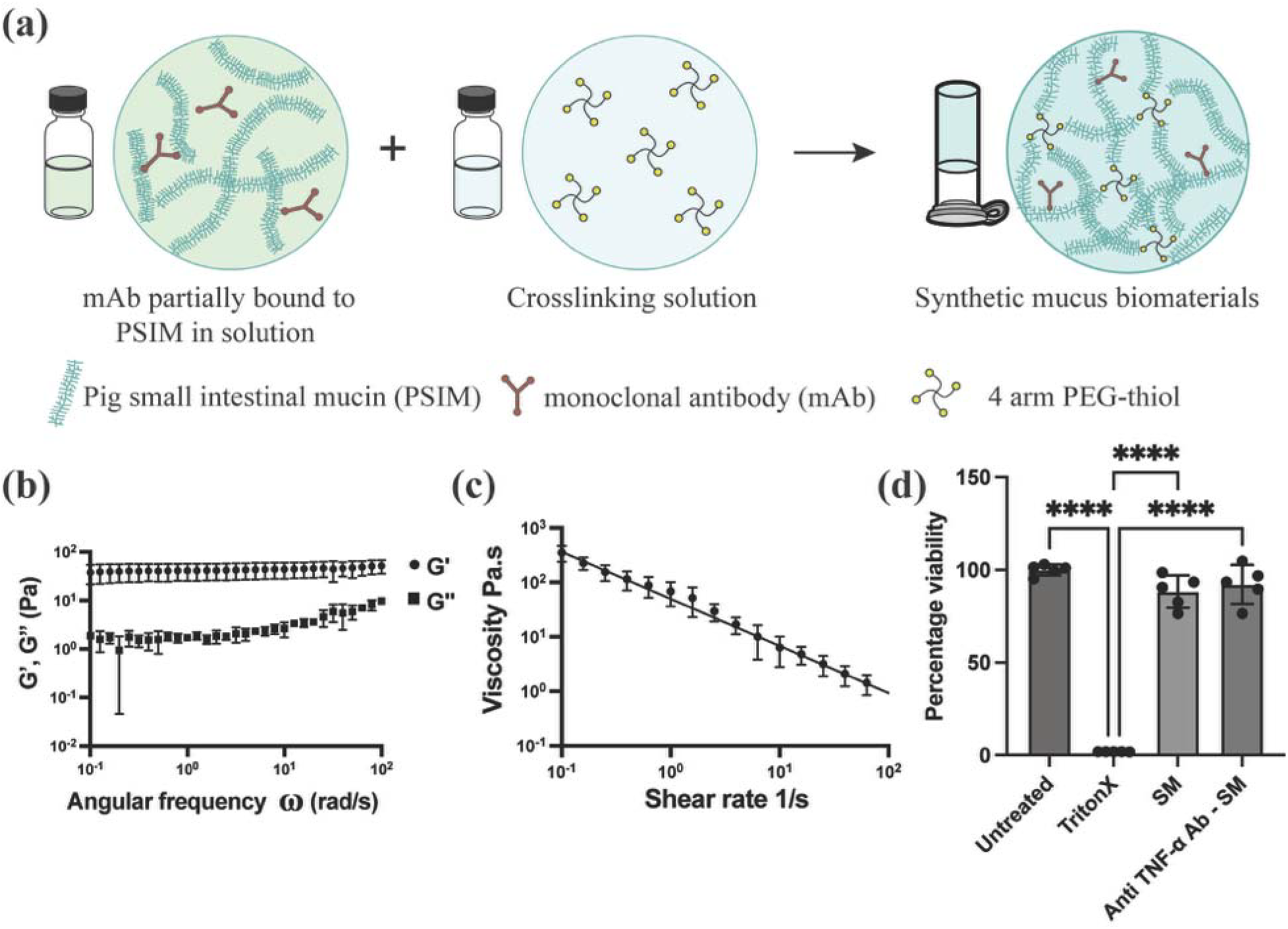
Synthesis and characterization of synthetic mucus gels for monoclonal antibody (mAb) delivery. (**a**) Schematic of SM gel formation through disulfide crosslinking and loading of mAbs through Fc mediated antibody interactions. (**b**) Storage (G’) and Loss (G”) modulus measure by frequency sweep at 1% strain. (**c**) Flow sweep showing decrease in dynamic viscosity with increase in shear rate. (**d**) Cell viability of HEK293T cells 24 hours post treatment with SM gels (n=5 technical replicates), \*\*\*\**p* <0.0001 for one-way ANOVA with Tukey’s multiple comparison test.

### Mucin–IgG interactions mediate controlled release of monoclonal antibodies from SM gels

Fluorescence recovery after photobleaching (FRAP) was employed to assess the mobility of IgG and its interactions with mucins within SM gels. The fluorescence recovery profile revealed a mobile fraction of 65% and an immobile fraction of 35%, indicating partial immobilization and/or reversible binding of IgG to the mucin backbone in the SM gels (**Fig. 2a**). As shown in **Fig. 2b**, the calculated diffusivity of IgG from the recovery half-time was 2.9 μm^2^/s, substantially lower than the diffusivity reported for IgG in native human mucus (∼29 μm^2^/s), reflecting a ∼10-fold reduction. This decrease in diffusivity is consistent with the reported binding of IgG to mucins via adapter molecules such as Fcγ binding protein (FcγBP) supporting controlled antibody release from the SM gels.

**Figure 2.**
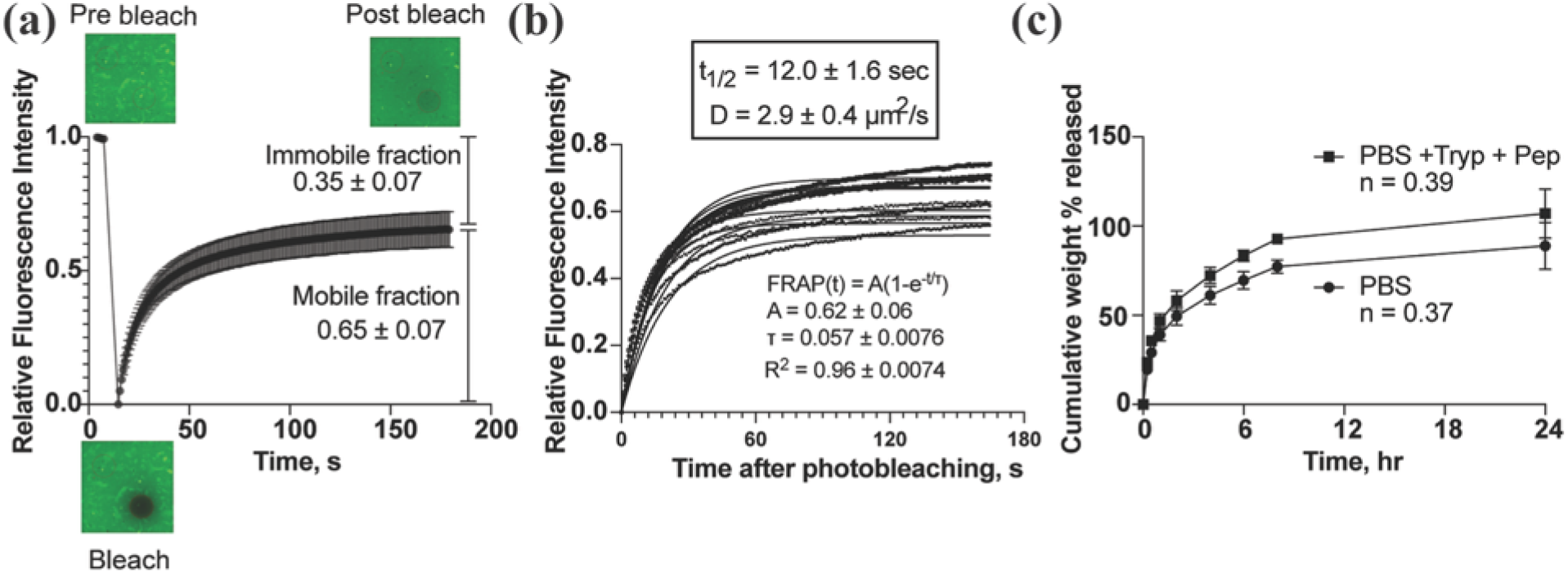
IgG antibody diffusion through and release from synthetic mucus gels. (**a**) Fluorescence recovery curve showing mobile and immobile fractions of IgG. (**b**) Diffusivity of IgG within synthetic mucus gel calculated from FRAP data fit to exponential model. (**c**) Cumulative IgG release from synthetic mucus gels in phosphate-buffered saline (PBS) and in PBS containing gastrointestinal proteases, representing conditions that mimic the intestinal environment (n = 3 biological replicates). Data shows meanL±LSD.

Drug release studies in PBS alone showed initial burst release of 20% and a total of 90% release within 24 hours. Drug release in conditions mimicking the protease rich gastrointestinal tract showed increased drug release reaching ∼100% in 24 hours. The enhanced release observed in the presence of proteases is attributed to the proteolytic degradation of the mucin network within the SM gels, which compromises the gel structure and facilitates drug diffusion. Analysis of the release kinetics using the Korsmeyer–Peppas model revealed a diffusion exponent (n) of < 0.45 in both PBS and protease conditions, indicating a Fickian diffusion-dominated release mechanism (**Fig. 2c**) for 3% PSIM + 2% PEG SM gels. However, for formulations containing higher mucin content such as 4% PSIM + 4% PEG SM gels, the release mechanism was altered in response to protease exposure. While the n value remained < 0.45 in PBS, it increased to > 0.45 in the presence of proteases, transitioning to anomalous transport, where both diffusion and matrix degradation contribute to drug release (**Fig. S2**). This shift highlights the role of IgG– mucin binding in drug release: under proteolytic conditions, degradation of the mucin backbone disrupts mucin–IgG interactions, thereby facilitating the release of the bound antibody.

### Modulation of macrophage activity by mucins and synthetic mucus gel

To assess macrophage function after treatment with synthetic mucus gels, we evaluated phagocytic activity of a murine macrophage cell line (RAW 264.7) using pHrodo-labeled *E. coli* bioparticles, which fluoresce upon internalization into the acidic endosomal compartment. Both fluorescence microscopy and flow cytometry confirmed active phagocytosis across all conditions. Unstimulated macrophages showed increased phagocytosis following SM treatment, with a higher proportion of pHrodo-positive cells compared to control (**Fig. 3a**). Morphologically, these SM-treated cells exhibited greater spreading, indicative of actin reorganization and associated with a pro-inflammatory phenotype. In LPS-stimulated macrophages, phagocytosis and cell spreading were observed under all conditions, and SM treatment did not significantly alter the percentage of pHrodo-positive cells compared to LPS alone (**Fig. 3b**). Analysis of mean fluorescence intensity (MFI) showed no significant differences between untreated and SM-treated cells within each treatment group, indicating that the extent of bioparticle uptake per cell remained unchanged. While LPS-stimulated macrophages exhibited higher MFIs overall, SM hydrogel treatment did not further increase this measure (**Fig. 3c**).

**Figure 3.**
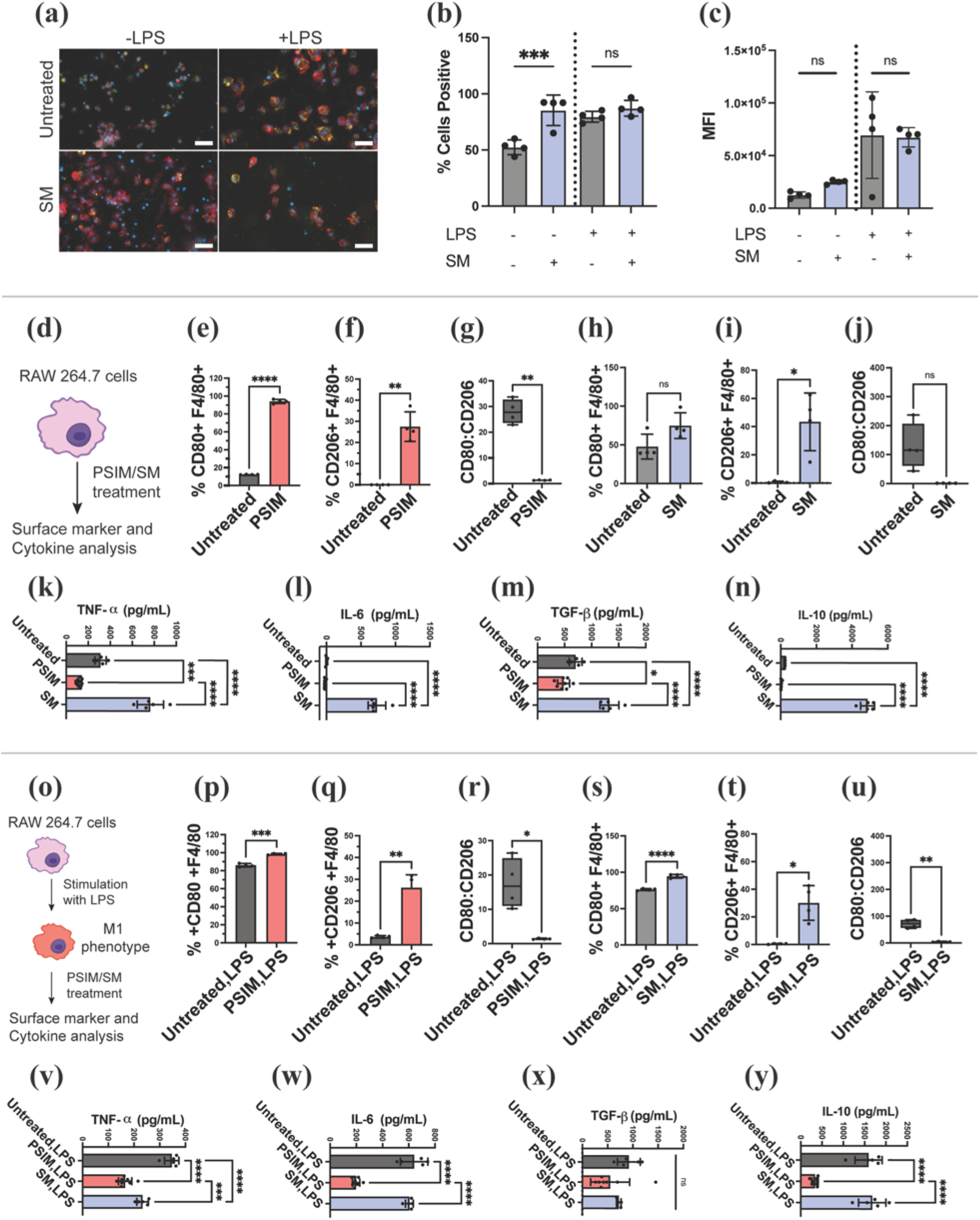
Immunomodulatory function of mucins (PSIM) and synthetic mucus (SM) biomaterials on macrophages. (**a**) Fluorescence imaging of pHrodo bioparticle uptake by RAW 264.7 macrophages. Macrophages were stained with phalloidin (red) and DAPI (blue) to visualize actin and nuclei, respectively (*n* = 4). Scale bar represents 50 µm. (**b**) The percentage of macrophages that phagocytized pHrodo bioparticles after treatment was quantified via flow cytometry (*n* = 4). (**c**) The mean fluorescence intensity (MFI) was measured using flow cytometry to determine the phagocytic capacity of macrophages that phagocytize pHrodo bioparticles (*n* = 4). non-significant (ns), \**P*<0.05, \*\*\**P*<0.001 and \*\*\*\**P*<0.0001 for one-way ANOVA. Data shows meanL±LSD. (**d**) Graphical depiction of study to examine impact of direct treatment with mucins in solution and hydrogel form on macrophage phenotype. (**e-j**) Flow cytometry analysis of cell surface markers, CD80, and CD206 in unstimulated macrophages (*n* = 4). (**k-n**) Cytokine production (TNF-α, IL-6, TGF-β, IL-10) in unstimulated macrophages determined by ELISA (*n* = 4). (**o**) Graphical depiction of study to examine impact of direct treatment with mucins in solution and hydrogels on LPS stimulated macrophage phenotype. (**p-u**) Flow cytometry analysis of cell surface markers, CD80, and CD206 in LPS stimulated macrophages (*n* = 5-8). (**v-y**) Cytokine production (TNF-α, IL-6, TGF-β, IL-10) in LPS stimulated macrophages determined by ELISA (*n* = 5-8). Data sets in (**e-j**) and (**p-u**) statistically analyzed via Welch’s t-test: \**P*<0.05, \*\**P*<0.01, \*\*\**P*<0.001 and \*\*\*\**P*<0.0001 for. Data sets in (**k-n**) and (**v-y**) statistically analyzed via one-way ANOVA: \**P*<0.05, \*\*\**P*<0.001 and \*\*\*\**P*<0.0001.

To further examine the immunomodulatory effects of mucins, either in solution or hydrogel form, we treated RAW 264.7 macrophages under both naive and LPS-stimulated inflammatory conditions to better resemble activated macrophage phenotypes in IBD. Upon exposure to PSIM or SM gels, macrophages exhibited polarization toward both pro-inflammatory M1-like (CD80) and anti-inflammatory, pro-repair M2-like (CD206) phenotypes with overall dominance of M2-like phenotype as evidenced by higher CD206/CD86 ratios (**Fig. 3d-j, Fig. 3o-u**). In addition, cytokine data showed a similar trend for unstimulated macrophages with increase in both pro-inflammatory (TNF-α and IL-6) and anti-inflammatory mediators (TGF-β and IL-10) for SM gel treatment (**Fig. 3k-n**). Under LPS-induced inflammatory conditions, gel-treated macrophages exhibited a reduction in pro-inflammatory cytokines, with minimal change in anti-inflammatory markers suggesting a net immune dampening effect with a shift away from the M1 (pro-inflammatory) phenotype without promoting M2 (anti-inflammatory) polarization (**Fig. 3v-y**). This highlights the robust and targeted immunomodulatory potential of SM hydrogels. In contrast, PSIM treatment caused a broad suppression of both pro-inflammatory (TNF-α and IL-6) and anti-inflammatory (TGF-β and IL-10) mediators in both unstimulated and LPS-stimulated macrophages (**Fig. 3k-n, Fig. 3v-y**). This generalized downregulation suggests a less selective, global dampening of macrophage cytokine production by PSIM. Taken together, these results suggest that SM hydrogel treatment enhances the recruitment of resting macrophages into a more active, phagocytic state potentially through interactions of Siglec receptors with mucin-associated glycans such as sialic acids.^17,18,25,26^ This could promote the resolution of inflammation by facilitating clearance of cellular debris and microbial products, a critical step in restoring tissue homeostasis. Overall, these findings indicate that mucin-based hydrogels can modulate macrophage phenotype and function while preserving essential immune functions such as phagocytosis. This immunoregulatory capacity has promising therapeutic implications for chronic inflammatory diseases like IBD, where impaired macrophage polarization and defective resolution of inflammation play a central role in disease progression.

### Anti-TNF_α_-loaded SM gels effectively neutralize TNF-_α_ and promote anti-inflammatory phenotype in LPS stimulated macrophages

RAW264.7 macrophages stimulated with LPS exhibited a pro-inflammatory response, with significant increase in TNF-α secretion compared to unstimulated controls. Treatment with blank SM gels did not alter TNF-α levels, indicating that the gel matrix alone does not interfere with TNF-α production. In contrast, anti-TNF-α antibody-loaded SM gels significantly suppressed TNF-α secretion compared to LPS treatment alone demonstrating the efficacy of the gel formulation in delivering functional neutralizing antibodies locally (**Fig. 4a**). Despite the significant reduction in TNF-α levels, IL-10 secretion remained unaffected across all LPS-stimulated conditions (**Fig. 4b**). These results suggest that IL-10 production in this context is TNF-α independent and may be driven alternatively by LPS-TLR4 signaling through MAPK/p38 and STAT3 activation pathways.^27–30^ Unlike systemic anti-TNF therapies, which have been shown to elevate IL-10 through Fcγ receptor engagement, this mechanism appears to operate independently of antibody-Fc receptor interactions.^31,32^ Further, analysis of macrophage surface markers revealed that treatment with both blank and anti-TNF-α-loaded gels significantly reduced CD80/CD206 ratio compared to LPS stimulated control indicating a shift toward an anti-inflammatory (M2-like) phenotype (**Fig. 4c**). No significant difference was observed between the two gel formulations, suggesting that the synthetic mucus matrix itself can drive macrophage polarization toward an M2-like phenotype independent of the anti-TNF-α payload. These findings are consistent with prior evidence showing that mucin-based hydrogels influence macrophage behavior through sialic acid mediated pathways, highlighting the potential of SM gels for localized immunomodulation and tissue repair.^17,18^

**Figure 4.**
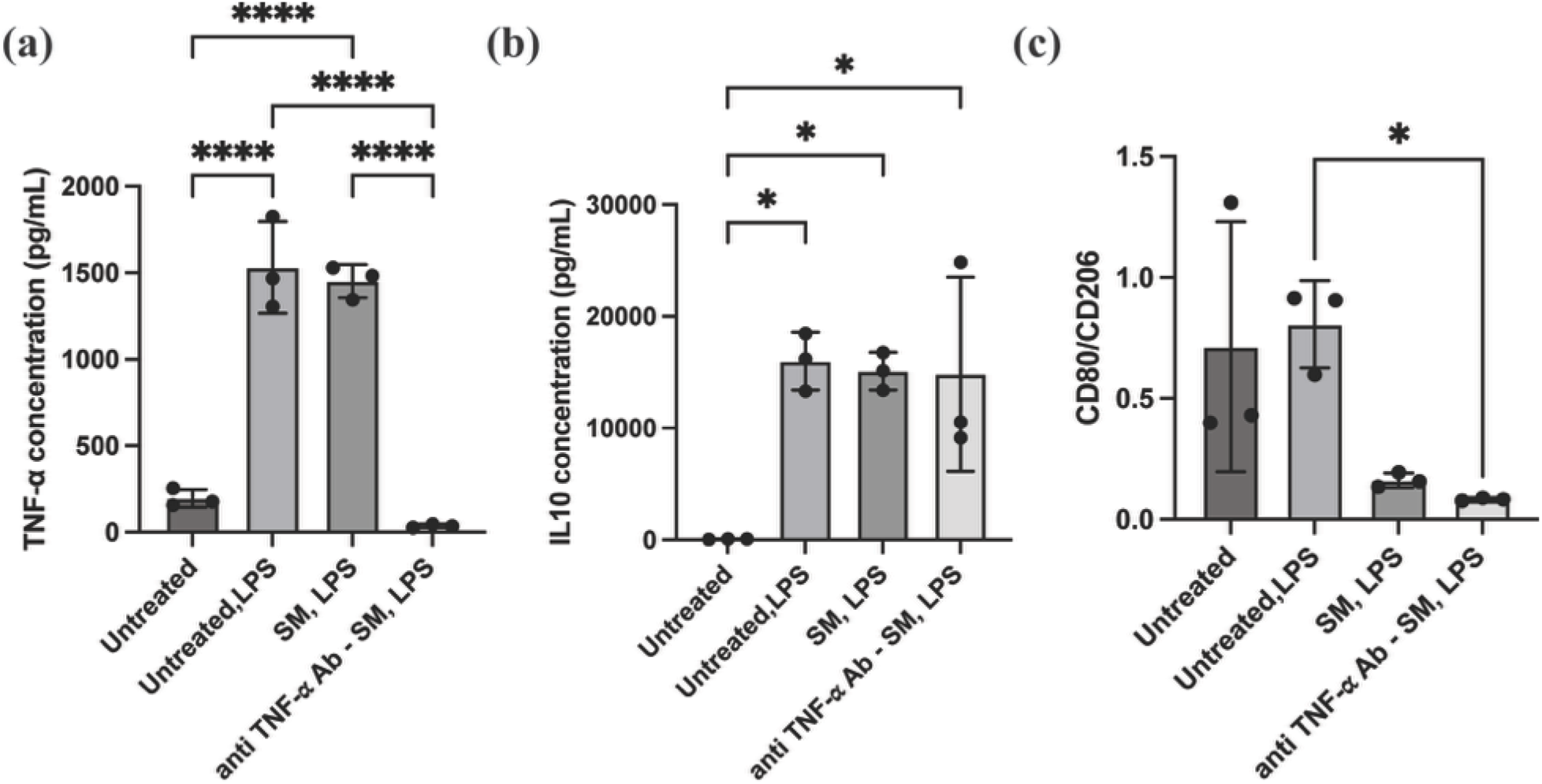
*In vitro* efficacy of Anti-TNF-_α_-loaded synthetic mucus gels. Production of (a) TNF-α, (b) IL-10, and (c) surface marker expression (CD80/CD206) in LPS stimulated RAW264.7 mouse macrophages following treatment with synthetic mucus gels (n=3 technical replicates), \**p* <0.05, \*\*\*\**p* <0.0001 for one-way ANOVA with Tukey’s multiple comparison test.

### Intrarectal antibody delivery using synthetic mucus gels

We used the TNBS-induced colitis model to evaluate the retention of IgG-loaded synthetic mucus gel formulation in comparison to IgG delivered in solution (**Fig. 5a**). In this model, TNBS dissolved in ethanol is instilled intrarectally to induce localized colonic inflammation, making it suitable for testing enema-based formulations for IBD. Following TNBS administration, we monitored body weight as an indicator of IBD induction. Mice exhibited a modest weight loss at 24 hours post-treatment, followed by gradual recovery (**Fig. 5b**). Notably, we did not observe other typical signs of colitis such as diarrhea, rectal bleeding, shortened colon, colonic wall thickening or increased tissue TNF-α concentration suggesting a mild inflammatory response (**Fig. S3**). Nevertheless, we proceeded with the retention study. To assess whether the SM gel improves mucosal retention, we intrarectally administered Cy5-labeled IgG either in SM gel or in solution. Since the IgG release occurs over a 24-hour period, the presence of fluorescence signal directly correlates with gel retention. As shown in **Fig. 5c** and **5d**, neither the solution nor the gel formulation was detectable at the 2-hour time point. However, when imaged shortly after administration (∼15 minutes), the SM gel group exhibited approximately two-fold higher fluorescence intensity compared to the IgG solution. These findings indicate that the synthetic mucus gel facilitates absorption of IgG at the colonic mucosa, supporting its potential as a local delivery platform for monoclonal antibodies.

**Figure 5.**
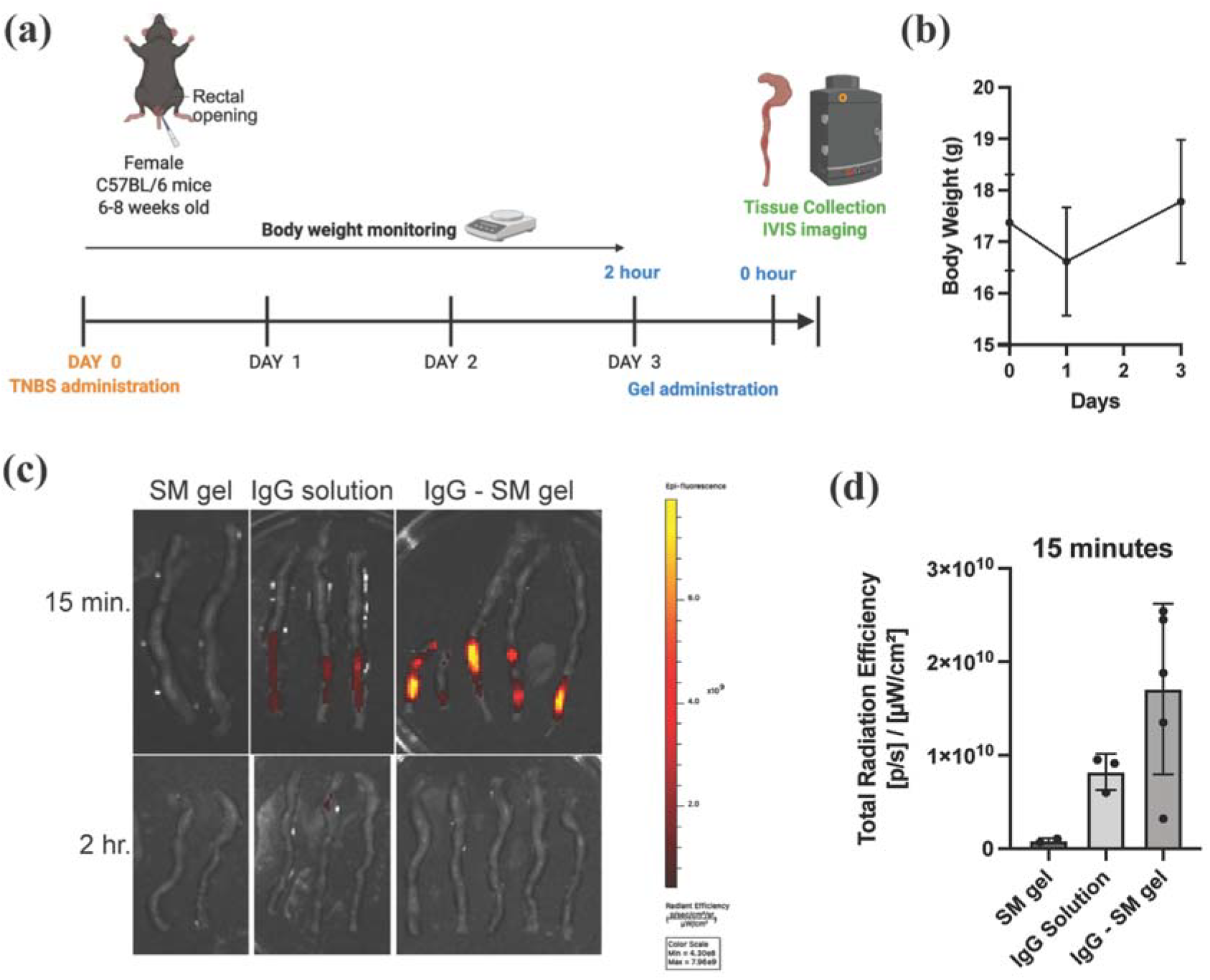
*In vivo* retention of synthetic mucus gels. (**a**) Study timeline for TNBS administration and intrarectal delivery of IgG and IgG-loaded synthetic mucus (SM) gel. (**b**) Body weight of mice monitored throughout the study to evaluate disease induction prior to treatment with antibody or gels. (**c**) *Ex vivo* IVIS images of mice colons following administration of either SM gel, Cy5 labeled IgG in solution, or Cy5 labeled IgG loaded in SM gel. (**d**) Quantification of total radiation efficiency from IVIS images, used to assess *in vivo* retention of IgG following administration. No statistical significance between groups for one-way ANOVA with Tukey’s multiple comparison test. Data shows meanL±LSD.

## Conclusion

Synthetic mucus gels present a promising platform for localized therapeutic immunomodulation at mucosal sites while minimizing systemic toxicity. We successfully formulated TNF-α targeting monoclonal antibody (mAb)-loaded synthetic mucus hydrogels demonstrating that these gels preserve mAb bioactivity and enable controlled, protease responsive drug release within the gastrointestinal tract. This approach addresses major limitations of conventional mAb delivery and can significantly enhance patient adherence. Furthermore, synthetic mucus gels can be used as drug-free therapeutics to directly lower inflammation, expanding their potential utility as immunomodulatory biomaterials in IBD and other inflammatory conditions.

## Acknowledgements

This work was supported by the NIH (R21 EB030834 awarded to GAD, KM; R01 HL160540 awarded to GAD) and NSF GRFP (GRFP DGE1840340 awarded to SY).

## Ethics Approval

All animal experiments were conducted in accordance with protocols approved by the Institutional Animal Care and Use Committee (IACUC protocol # R-NOV-22-50) at the University of Maryland.

## Author Contributions

**Conceptualization:** TY, SY, KM, GAD

**Methodology:** TY, SY, MK, RL, SD, KM, GAD

**Investigation:** TY, SY, MK, RL, SD

**Visualization:** TY, SY, GAD

**Supervision:** KM, GAD

**Writing—original draft:** TY, SY, GAD

**Writing—review & editing:** TY, SY, MK, RL, SD, KM, GAD

## Data Availability

The data that supports the findings of this study are available from the corresponding author upon reasonable request.

