## Supplementary Information for "Synthetic Mucus Biomaterials Enable Localized Therapeutic Antibody Delivery in Inflammatory Bowel Disease"

**Table S1**. Degradation times of different formulations of SM gels.

| Formulation number | PSIM (%w/v) | 4-arm PEG-SH  (%w/v) | Gel formation | Degradation time in PBS |
| --- | --- | --- | --- | --- |
| 1 | 4 | 4 | Yes | 10 days |
| 2 | 4 | 2 | Yes | 4 days |
| 3 | 4 | 1 | Yes | < 24 hours |
| 4 | 4 | 0.5 | No | - |
| 5 | 3 | 1 | No | - |
| 6 | 3 | 0.5 | No | - |
| 7 | 3 | 2 | Yes | > 24 hours |
| 8 | 2 | 2 | Yes | 6 hours |

**Video S1. Injectability of SM Gel Through an 18G Needle.**
This video demonstrates the injectability of the SM gel formulation through an 18-gauge needle. As shown, increasing the concentration of PSIM (labeled as PGM in the video) results in greater difficulty during injection, indicating a concentration-dependent increase in resistance to flow.

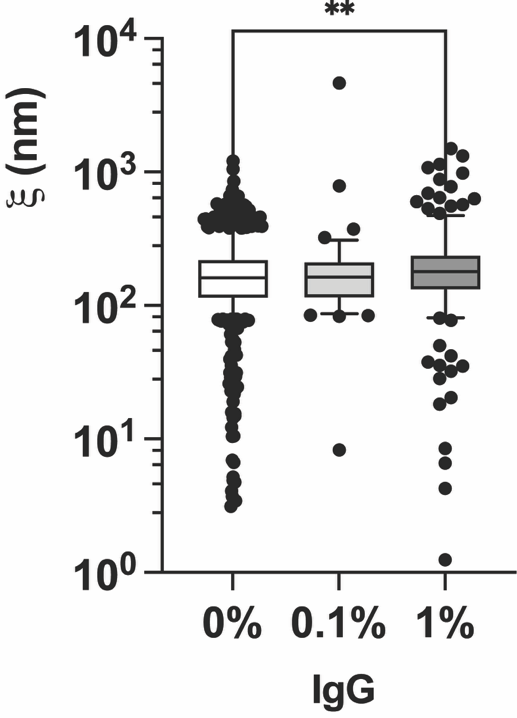

**Figure S1. Microrheological properties of** **3%PGM + 2%PEG gels.** Estimated pore size for particle tracking microrheology using densely PEGylated 100 nm nanoparticles as probes. * *p* <0.05, ****p* <0.001, *****p* <0.0001 for Kruskal-Wallis test.

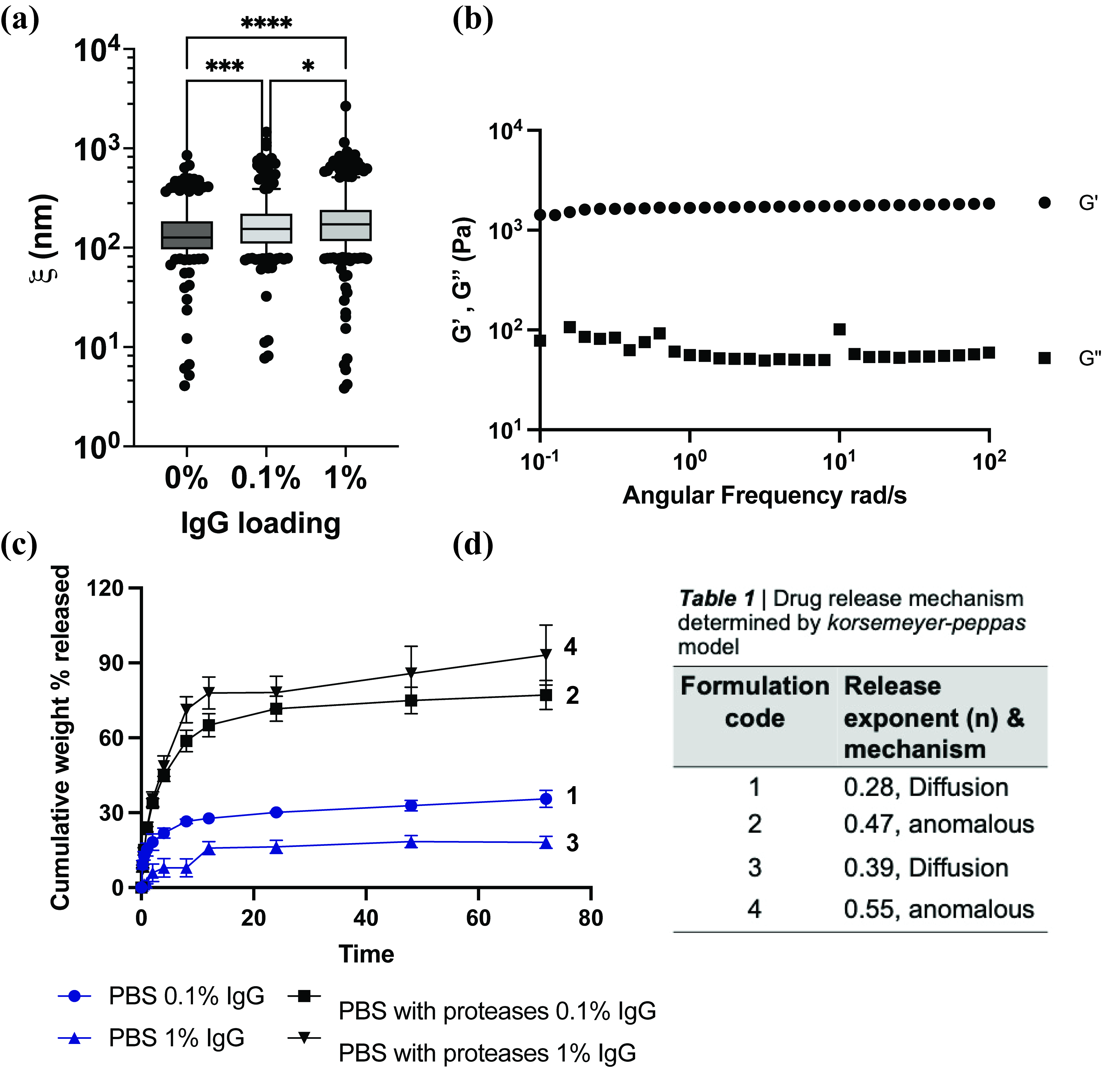

**Figure S2.** **Physical characterization of and drug release from of 4%PGM + 4%PEG.** (a) Estimated pore size size for particle tracking microrheology using densely PEGylated 100 nm nanoparticles as probes. * *p* <0.05, ****p* <0.001, *****p* <0.0001 for Kruskal-Wallis test. (b) Storage modulus (G’) and Loss modulus (G”) as a function of frequency at 10% strain. (c) Cumulative IgG release from synthetic mucus gels in phosphate-buffered saline (PBS) and in PBS containing gastrointestinal proteases, representing conditions that mimic the intestinal environment (n = 3 biological replicates). (d) Table with release exponent and mechanism of drug release. Data shows mean ± SD.

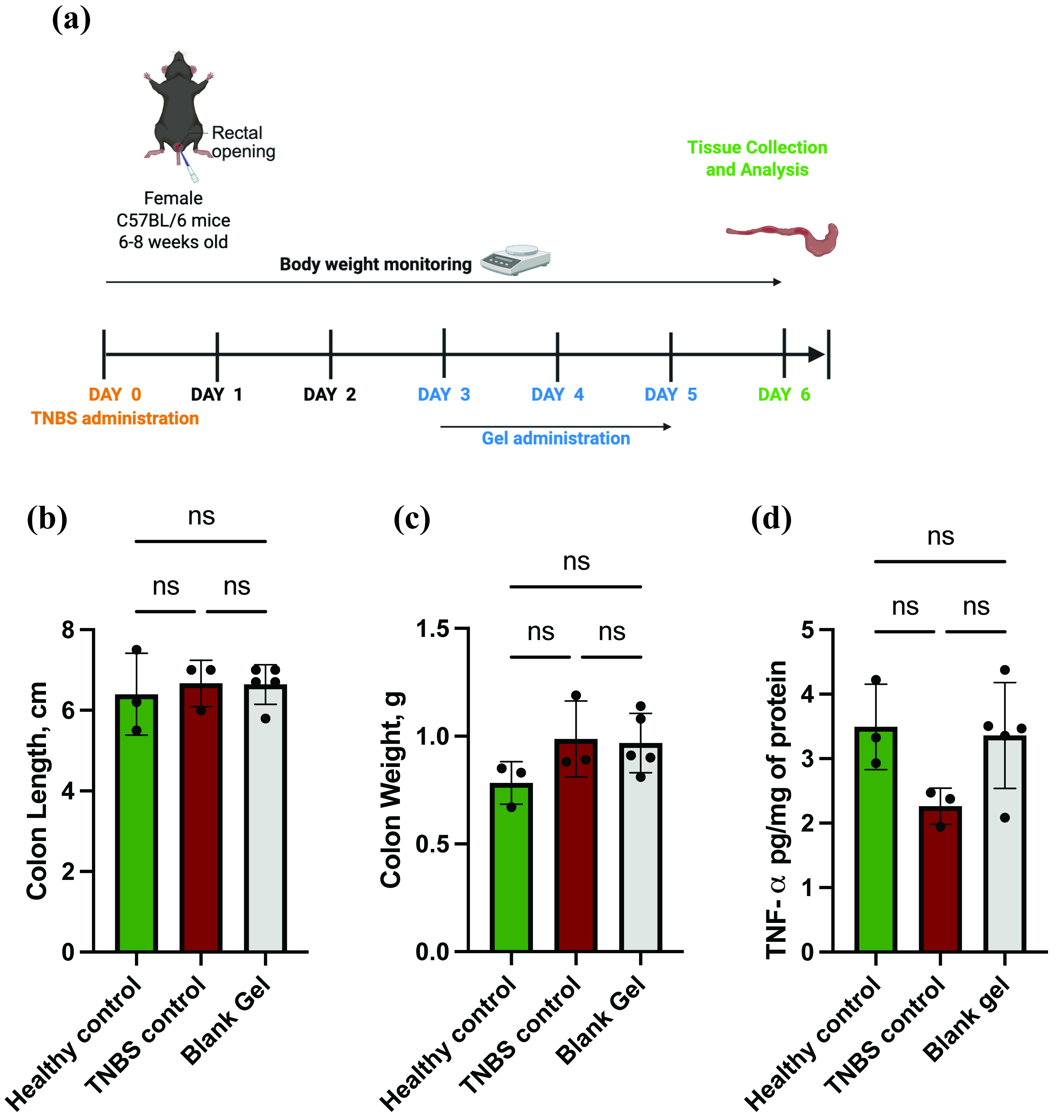

**Figure S3. In vivo efficacy of synthetic mucus gels.** (a) Study timeline for TNBS administration and intrarectal delivery of synthetic mucus (SM) gel. (b) Colon length, (c) Weight of distal 5 cm of the colon, and (d) Tissue TNF-α levels of healthy control (n=3), TNBS control (n=3), and SM gel treated group (n=5) at the end of the study. No statistical significance between groups for one-way ANOVA with Tukey’s multiple comparison test. Data shows mean ± SD.
